# Human prion proteins and mutants through the lens of systems biology

**DOI:** 10.64898/2026.10.24.761545

**Authors:** Swati Sharma, Faruque Siddique, Deep Nath, Jayanta Mukhopadhyay, Soumen Roy

## Abstract

Autocatalytic conformational changes of cellular prion proteins (PrPᶜ) to the β-sheet–enriched scrapie isoform (PrP^sc^) drive prion diseases, yet the precise molecular mechanisms remain unclear. To address this, we apply a multi-pronged framework integrating network-theoretic analyses, molecular dynamics (MD) simulations, and experimental validation to characterize critical residues governing human PrP (huPrP) structural integrity. Residue Interaction Networks (RINs) of: (A) wild-type (WT) huPrP, and, (B) five disease-associated point mutants (Met166Val, Glu220Lys, Glu200Lys, Asp178Asn, and Ser170Asn) were evaluated using three rigorous network methods: centrality measures, **<u>S</u>**hortest **<u>P</u>**ath **<u>A</u>**lteration **<u>F</u>**raction (SPAF), and **<u>D</u>**ifferential **<u>N</u>**etworks (DN). Across all the diverse point mutations, five residues (Met206, Val209, Val210, Met213, and Cys214) consistently exhibit high centrality scores. SPAF and DN analyses demonstrate that alterations at these positions disrupt connectivity and increase the propensity for misfolding and protease resistance. To validate these topological findings, key residues within the C-terminal domain (121-230) were substituted via site-directed mutagenesis. Thermal denaturation monitored by circular dichroism (CD) spectroscopy reveals that the Met213Ala substitution significantly lowers the melting temperature, T_m_, relative to the WT, whereas the Val210Ala variant unfolds with a T_m_ comparable to the WT. Furthermore, MD simulations of de novo AlphaFold-predicted mutant structures corroborate these findings. Mutants exhibit greater structural flexibility and deviation, with average root Mean Square Deviations rising from 0.18 nm (WT) to 0.32 nm (Cys214Ala), 0.29 nm (Met206Ala), and 0.25 nm (Val210Lys). This integration of theoretical and experimental techniques provides a robust framework for identifying critical determinants of protein stability.

## Introduction

Proteins are complex chains of amino acids joined by peptide bonds, particularly in a unique sequence. The chain folds itself and makes a 3D structure. This folding process is much more than a mere random overlapping and is highly critical because it enables the protein to execute its function efficiently. However, folding errors can lead to disastrous diseases such as prion diseases. Prion diseases are fatal transmissible neurodegenerative disorders caused by the misfolding of the cellular prion protein (PrP^C^). The conversion of PrP^C^, which acts as a cell surface glycoprotein (1), into a misfolded scrapie pathogenic protein (PrP^Sc^) leads to the formation of an aggregation-prone isoform (2). The misfolded isoform of the prion protein accumulates in an amyloid-like structure and targets the signal transduction (3) in the central nervous system of individuals (4).

The mechanism of conversion from the soluble monomeric form of PrP^C^ to the insoluble aggregated PrP^Sc^ form is still unclear. Among all the hypotheses proposed for the cause of prion diseases, the ‘protein-only’ hypothesis is the most widely accepted, supported by experimental (5) evidence that suggests the disease is caused entirely by the protein itself (6). Once PrP^Sc^ is formed, it acts as a template that promotes further amyloid accumulation by converting PrP^C^ into additional PrP^Sc^.

Prion diseases can be introduced into the human body by two routes, external sources (acquired) and internal sources (genetic or sporadic). Point mutations in the PRNP gene (which encodes PrP) are the primary reason for the accumulation of scrapie form through internal sources (7).

Structurally, after post-translational cleavage, the mature human cellular prion protein (huPrP^C^) consists of 23-230 amino acids and comprises two distinct domains: an N-terminal disordered region (23-120) and a C-terminal globular domain. The three-dimensional structures of huPrP^C^ have been extensively studied by using nuclear magnetic resonance and X-ray crystallography (1, 8–10). The C-terminal domain consists of three *α* helices and two antiparallel *β* sheets that span residues 144-154, 173-194, 200-228, 128-131, and 161-164, respectively, and a disulphide bond between Cys179-Cys214 (11).

Single-nucleotide mutations in the PRNP gene are found to increase the likelihood of misfolding and neurodegeneration (3). Numerous pathogenic mutations in the human prion protein gene (PRNP) have been identified and are linked to inherited prion diseases, including familial Creutzfeldt-Jakob disease (fCJD), fatal familial insomnia (FFI), and Gerstmann-Sträussler-Scheinker syndrome (GSS) (3, 12).

Herein, we incorporate a rigorous graph-theoretical approach to study the topological alteration in the structure of huPrP and its mutants (Met166Val, Ser170Asn, Glu200Lys, Arg220Lys, and Asp178Asn) using residue interaction networks (RIN). Where nodes represent the amino acids and edges represent the inter-residue contacts within 4.5Å (13). After employing topological centrality measures, we used **<u>D</u>**ifferential **<u>N</u>**etworks (DN) (13) and **<u>S</u>**hortest **<u>P</u>**ath **<u>A</u>**lteration **<u>F</u>**raction (SPAF) (14), based approaches. Remarkably, DN and SPAF also lead to the identification of highly significant residues that were shown to be important from the topological analyses of the residue interaction networks of the proteins. Within the networks of both WT and mutant proteins, a subset of residues (Met206, Val209, Val210, Met213, Cys214) shows high topological importance, with functional and structural relevance, and could be essential for maintaining the prion protein in its cellular isoform. We hypothesize that a mutation in these functional residues may alter the intramolecular network of huPrP^C^, thereby increasing the propensity for PrP^C^ to convert to PrP^Sc^.

Structural properties of WT huPrP (121-230), Val210Ala and Met213Ala variants were studied by CD-monitored thermal denaturation. We compared the conformational stability of WT huPrP(121-230) with that of the Val210Ala and Met213Ala variants using far-UV circular dichroism and thermal denaturation in combination with enhanced Molecular Dynamics simulations. We analyze the backbone RMSD and per-residue RMSF to assess how each substitution alters the structure and its interactions. Thus, our work seeks to bridge topological network measures and pathogenic outcomes in prion diseases, providing a route based upon predictions arising from topological analyses of the RIN to testable structural hypotheses.

## Materials and Methods

### 1. Residue Interaction Networks

An extensive analysis of huPrP and its potent mutants was conducted using graph theoretic approaches. Experimentally determined structures were retrieved from the Protein Data Bank (15, 16): 1qm2 (WT), 1e1g (Met166Val), 1e1p (Ser170Asn), 1fkc (Glu200Lys), 1e1w (Arg220Lys) and 2k1d (Asp178Asn). For each structure, an unweighted RIN is constructed, in which nodes represent amino acid residues, and edges represent inter-residue contacts. We consider not only the distant influence of amino acids but also the influence of adjacent/backbone amino acids during the construction of the network.

Protein structures can be represented as a graphs *G* = (*V*, *E*), *V* is the set of nodes representing amino acids and *E* is the set of edges between the *i^t^*^ℎ^ and *j^t^*^ℎ^residue, provided, the *i^t^*^ℎ^ and *j^t^*^ℎ^nodes are such that:

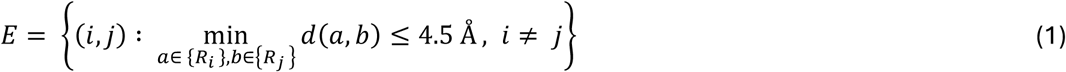

Here, {*R_i_*} and {*R_j_*} are the set of atoms belonging to the residues, *i*, and, *j*, respectively. Further, *d*(*a*, *b*) is the Euclidean distance between atoms *a* ∈ {*R_i_*}, and, *b* ∈ {*R_j_*}.

We construct an edge between the *i^t^*^ℎ^ and *j^t^*^ℎ^ residue if the distance between any atom, *i* ∈ {*R_i_*}, and, any atom *j* ∈ {*R_j_*}, ∀*_i_*_,*j*_ is ≤ 4.5 Å (17). Hence, the RIN represents the structural connectivity map of residues based on the atomic proximity between residues (18, 19).

It should be especially noted that, throughout herein, we will use the words “measure” and “metric” interchangeably, in the context of topological analyses of networks.

#### A. Topological analyses of Residue Interaction Networks

##### Betweenness centrality

The betweenness centrality, *B_m_*, of a node, *m*, indicates its role as a “topological bridge” within a network *G* and is defined as:

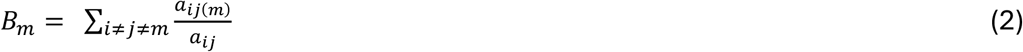

*a_ij_*(*m*) Is the number of shortest paths between *i^t^*^ℎ^ and *j^t^*^ℎ^ nodes that pass through the node *m*, while, *a_ij_*, is the total number of shortest paths between the *i^t^*^ℎ^ and *j^t^*^ℎ^ nodes in the network (18, 19). It has been reported that applying this measure to RINs identifies amino acids that govern the energy and information transfer through the protein. Residues with higher *B* are typically conserved functional hotspots that dictate allosteric pathways and structural stability; mutations at these specific nodes often lead to severe destabilization or misfolding (17, 18).

##### Closeness centrality

*C_i_*, of a node *i*, measures the inverse of the average shortest path length from the node *i*, to every other node, *j*, present in the network. Thus,

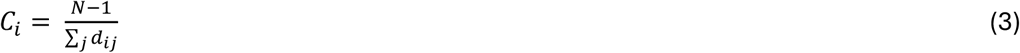

where *N* denotes the total number of nodes and *d_ij_*, denotes the length of the shortest path from the *i^t^*^ℎ^ node to *j^t^*^ℎ^ node (20). Generally, closeness centrality indicates how

“close” a node is to other nodes. A higher value of closeness implies that the node is centrally located and can interact quickly with the rest of the network. In RINs, residues with a high C_i_ readily connect to the rest of the protein and often serve as important functional active sites (13, 21, 22).

##### Eigenvector centrality

Eigenvector centrality, *E*, typically indicates the importance of nodes within the network. The calculation of *E* requires the principal eigenvector, i.e., eigenvector corresponding to the eigenvalue with highest absolute value, *λ*, of the associated adjacency matrix, *A* = [*a_ij_*]. Unlike degree centrality, *E* considers the influence of neighboring nodes by assigning a score depending on the values of elements of the principal eigenvector of *A*, i.e., *X* = [*x_j_*] (23–25). The eigenvector centrality of *i^t^*^ℎ^ node is calculated as,

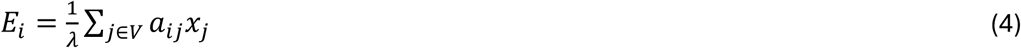

In RINs, *E_i_*, is useful in finding allosteric communication pathways and identifying key amino acid residues responsible for transferring structural information (26). Furthermore, it is highly sensitive to subtle structural shifts, making it uniquely capable of distinguishing salient structural features between different protein conformational states (27).

##### Information centrality

While calculating betweenness and closeness centrality, we considered the shortest paths. However, information centrality, *I*, considers all possible paths between two nodes weighted based on information-based values. Typically, *I_i_* calculates the harmonic mean length associated with all paths ending at *i^t^*^ℎ^ node. Therefore, *I_i_* helps in understanding how effectively information flows within a network, providing a more comprehensive assessment of *i^t^*^ℎ^ node’s importance (28). Mathematically,

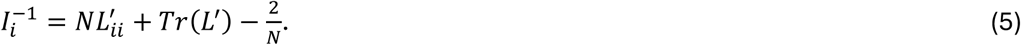

Here, *L*^′^ = (*L* + *J*)^−1^. *L* represents the Laplacian of the underlying graph and *J* = 11*^T^*. It has been reported that *I_i_* can also be obtained by evaluating the effective resistance between nodes, where each edge is treated as a resistor. In the context of resistor networks, *I* is often interpreted as the “current-flow closeness centrality” (29),

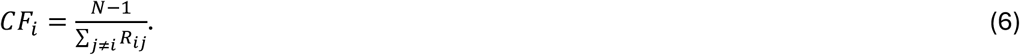

Here, effective resistance, *R_ij_*, between nodes *i* and *j* can be evaluated from the Moore-Penrose pseudoinverse of *L*. Biologically, this measure maps all possible communication pathways to effectively evaluate the overall structural stability and compatibility of interacting protein subunits (18).

There are various other useful network measures like edge proximity, *etc* (28, 29). However, not all network measures are useful in any given scenario, and there is a body of work in the literature to assess the suitability and appropriateness of any measure in a given scenario (31, 32).

##### Rescaling of Metric values

Normalization was applied to visualize the cumulative impact of individual metric values (betweenness centrality, closeness centrality, information centrality and eigenvector centrality). The rescaled metric value for *i^t^*^ℎ^ node was computed as,

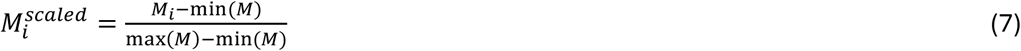

The scaled values of all the measures were then averaged for each node, and finally, nodes with average values exceeding 1.25 standard deviations above the mean were considered (18).

#### B. Shortest path alteration fraction calculations

The calculation of SPAF, *F^l^i*_,*j*,*li*,*j*_′ between two shortest paths, *l_i_*_,*j*_and *l_i_*_,*j*_, between *i* and *j* node present in both networks, *G* and *G*′, is defined as,

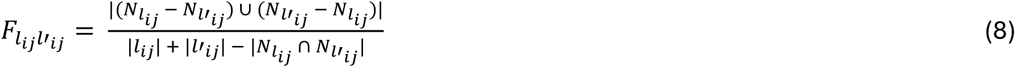

In the above equation, *N_lij_* and *N_ij_*^′^ represent the set of nodes lying on the shortest paths denoted by *l_i_*_,*j*_ and *l*^′^ in the network *G* and *G*′, respectively. Here, *G*′ was created upon removal of the *x^t^*^ℎ^ node from *G*. A value of *SPAF* = 1 represents completely altered paths between two given nodes.

The total number of shortest path pairs between *i^t^*^ℎ^ and *j^t^*^ℎ^ nodes in *G* and *G*′ for which SPAF=1 can be computed as:

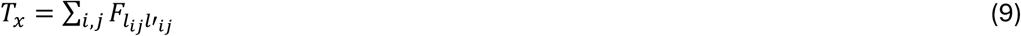

*T_x_*, is the total number of altered paths after removal of *x^t^*^ℎ^ node from *G*.

#### C. Differential Networks

The approach of differential networks (DN) specifically highlights the important interactions and the nodes involved in unique interactions in WT PrP^c^ (PDB: 1qm2) and mutants ((1e1g (Met166Val), 1e1p (Ser170Asn), 1fkc (Glu200Lys), 1e1w (Arg220Lys) and 2k1d (Asp178Asn)) separately. The DN is created by comparing the mutant and WT networks. The edges that are present in the mutant network but absent in the WT network, form the DN of the mutant protein (13). Conversely, the edges that exist in the WT network but are missing in the mutant network, constitute the DN of the WT protein.

##### Change in the value of a network metric

The change in the value of a network metric (*M*) such as (a) betweenness centrality, and (b) closeness centrality, of *i^t^*^ℎ^ node from average metric values (respectively) upon removal of *k^t^*^ℎ^ edge (removal of *k^t^*^ℎ^ edge will create (*G*′′)) from the parent network (*G*) (as well as from mutant states) can be mathematically represented as,

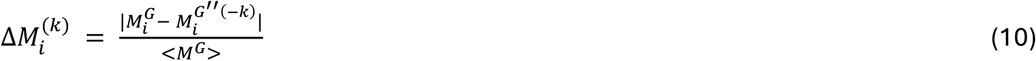

*M_i_^G^* is the metric value of node *i* in the original graph (*G*), and *M_i_^G’’(-k)^* is the metric value of graph (*G*′′) upon removal of *k^t^*^ℎ^ edge. < *M* > is the average of the metric values of the parent graph (*G*), which could be the WT and mutant-type network.

Furthermore, the overall impact of removing the *k^t^*^ℎ^ edge on the network’s metric can be quantified as,

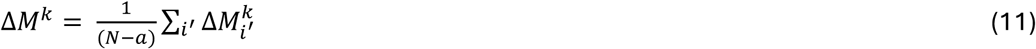

Here, *i*^′^ is the number of nodes excluding the incident nodes of the *k^t^*^ℎ^edge (13).

### 2. ΔΔG prediction

Changes in folding free energy (ΔΔG) upon point mutation were predicted using the CUPSAT server (33), which evaluates the mutation site using amino acid–atom pair potentials and main-chain torsion-angle distributions. The WT structure (PDB: 1qm2) was submitted and the mutation of interest specified under the thermal-stability option. Predicted ΔΔG values were interpreted according to the server’s convention. Positive values indicating a stabilising and negative values a destabilising effect relative to the WT.

### 3. Cloning and Protein expression

#### A. Synthesis of gene

A synthetic gene encoding residues 121-230, the structured C-terminal domain (8, 34) of the huPrP (121-230) [UniProt P04156, Met129 variant], codon-optimized for expression in *Escherichia coli*, was cloned between the BamHI and XhoI sites of pET-28a(+) downstream of the T7 promoter (35) and in frame with the vector-encoded N-terminal hexahistidine tag.

#### B. Site-directed mutagenesis

Mutations were introduced individually into the WT huPrP (121-230)/pET-28a(+) construct by whole-plasmid site-directed mutagenesis using complementary mutagenic primer pairs. PCR reactions contained 1× Phusion GC master mix, 150–200 ng template, 0.5 µM each primer and 3% (v/v) DMSO. The methylated parental template was removed by DpnI digestion (5 U, New England Biolabs), and the product was transformed into chemically competent *E. coli* DH5α with selection on LB agar containing 50 µg/mL kanamycin. Plasmid DNA from individual colonies was verified by Sanger sequencing using T7 promoter and T7 terminator primers (36).

#### C. Expression and purification

Verified constructs were subsequently transformed into BL21 (DE3) for expression, and mutant proteins were expressed and purified in parallel with the WT using identical conditions. Cell pellets were resuspended in 5 mL (per 50 mL of culture) of ice-cold lysis buffer comprising 50 mM Tris-HCl (pH 8.0), 300 mM NaCl and 10mM imidazole, supplemented immediately before use with 1mM PMSF, and 0.1mg/mL lysozyme. Suspensions were incubated on ice for 15-20 min with gentle rocking and lysed by sonication on ice (4 × 30 s bursts at 50% amplitude, with 30 s cooling intervals). Lysates were clarified at 13,000 × g for 20 min at 4 °C, and SDS-PAGE analysis of both fractions confirmed that the recombinant protein partitioned almost entirely into the insoluble pellet as inclusion bodies. Inclusion bodies were washed three times by resuspension in 3 mL of 50 mM Tris-HCl pH 8.0, 300 mM NaCl, 20 mM imidazole, followed by centrifugation at 12,000 × g for 10 min at 4 °C. Washed pellets were solubilised in 3 mL of denaturing buffer (6 M GndHCl, 50 mM Tris-HCl (pH 8.0), 300 mM NaCl, 10 mM imidazole) for 2 hr at room temperature with gentle rotation, and insoluble material was removed by centrifugation at 12,000 × g for 20 min at 4 °C. The solubilised protein was applied to 1mL of Ni–NTA agarose resin pre-equilibrated in the same denaturing buffer and incubated for 1 h at room temperature with end-over-end rotation. The resin was washed with 15 mL of denaturing wash buffer (as above, with 50 mM imidazole), and bound protein was eluted in 1mL fractions with elution buffer (6M GndHCl, 50 mM Tris-HCl pH 8.0, 300 mM NaCl, 500 mM imidazole). Fractions were analysed by SDS-PAGE and those containing the target protein were pooled. Refolding was achieved by dialysis (34, 37). Pooled eluate was transferred to dialysis tubing (MWCO 6–8 kDa) and dialysed at 4 °C against refolding buffer (20 mM Tris-HCl (pH 9.0), 150 mM NaCl); any precipitate was removed by centrifugation at 12,000 × g for 10 min. The clarified supernatant was then concentrated by ammonium sulphate fractionation: solid (NH₄)₂SO₄ was added slowly with gentle stirring at 4 °C to 60% saturation, and after 30 min of equilibration, the suspension was centrifuged at 12,000 × g for 20 min at 4 °C. Purified protein pellets are stored at – 80 °C.

### 4. CD spectroscopy

Circular dichroism spectra were recorded on a Jasco spectropolarimeter using a thermostatted quartz cuvette of 1 mm path length. Far-UV spectra were acquired over 190–260 nm in 10 mM sodium phosphate buffer, pH 7.0. Thermal denaturation was followed by monitoring the ellipticity at 222 nm from 30 to 90 °C in 5 °C increments, with the sample equilibrated at each temperature before acquisition. Melting curves were normalised to fractional unfolding, and the melting temperature (T_m_) was taken as the midpoint of the transition.

### 5. Molecular dynamics simulations

NMR structure of WT huPrP (PDB: 1qm2) and mutants (gly126ala, His140ala, Met206lys, Val209lys, Val210ala, Val210lys, Met213ala, and Cys214ala) were used as an initial structure for simulations. Three-dimensional structural coordinates of mutants were predicted by *de novo* modelling using AlphaFold 3. For each mutant, the top-ranked model was selected based on its high confidence score (mean pLDDT[X]). Atomic coordinates of the WT structure were retrieved from the RCSB Protein Data Bank and simulated independently as a reference.

We perform MD simulations with GROMACS 2026.3 (38) using the CHARMM36m force field (39)and the CHARMM-modified TIP3P water model (40). Each structure was placed in a rhombic dodecahedral box with a minimum distance of 1.2 nm between the protein and the box edge, solvated, neutralised, and brought to 0.15 M NaCl. The systems were energy-minimised by the steepest descent, then equilibrated for 100 ps in the NVT ensemble at 300 K and for 100 ps in the NPT ensemble at 1 bar, with position restraints on the protein heavy atoms. Production MD runs were carried out without any restraints at a constant temperature (300 K) and pressure (1 bar), using the velocity-rescaling thermostat (41) and the [C-rescale/Parrinello–Rahman] barostat (42). For each structure, three independent simulations were performed with different initial velocities. Each simulation was carried out for 500 ns using a 4 fs time step, yielding 30 simulations and a total of 15 µs of simulation time.

We analysed MD trajectories with GROMACS tools and MDAnalysis and python (43). From each trajectory, the first 50 ns were discarded as equilibration, and analysis was performed on the remaining 450 ns, with frames saved every 10 ps. The backbone and Cα root-mean-square deviation (RMSD) were calculated relative to the starting structure after least-squares fitting of each frame onto the backbone atoms of the whole protein. The per-residue Cα root-mean-square fluctuation (RMSF) was calculated about the average structure after the same fitting.

## Results

### Quantifying network topology by calculating centrality distributions of WT and mutant prion proteins

To enhance our understanding of prion protein structures, a comparative study of prion protein and its mutants was carried out. First, we analyse the distribution profile of network measures to assess structural changes in PrP^c^ before and after point mutations (WT, PDB: 1qm2). In Figure 1, the difference in distributions of closeness centrality and information centrality, along with other centrality measures, illustrate how mutations influence the structure of the network. The closeness centrality distribution of 1e1w resembles that of the WT (PDB: 1qm2), whereas 1fkc shows greater deviation. This indicates that the huPrP (Arg220Lys) mutation has minimal impact on residue connectivity, while the huPrP (Glu200Lys) mutation results in significant structural alterations. Some residues gain a higher value of centrality, enhancing inter-residue communication, while others lose connectivity, disrupting long-range interactions. A similar pattern is observed in other mutants (1e1g, 1e1p and 2k1d) where distributions remain largely consistent with the WT. However, in 1e1w, the differences are more pronounced, suggesting structural changes that directly affect edge connections. Since the frequency distributions show a high degree of similarity, we employ the Wasserstein divergence to accurately quantify the node-wise differences in centrality measures. A higher W-value indicates lower similarity to the WT, while a lower W-value reflects a higher degree of similarity.

**Fig 1:**
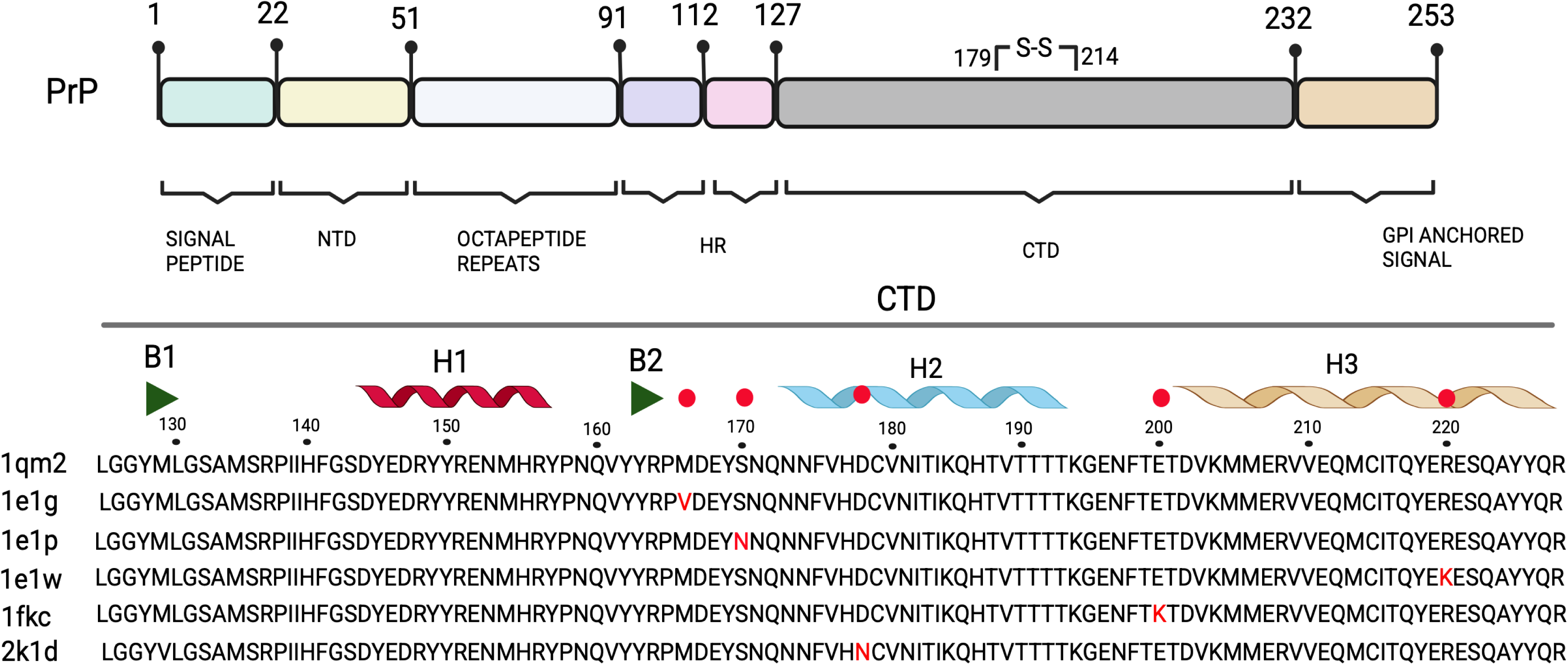
Outline of the full-length structure of the hPrP WT and point mutants. Alignment of the C-terminal globular domain (CTD) of WT and mutant proteins from 125-228 amino acids. CTD contains three alpha helices, H1, H2, and H3, and two beta sheets, B1 and B2. At the top of the alignment, secondary structure elements are indicated. Highlighted red letters identify the residue positions of substitution point mutations in the region within the CTD, as mentioned in the paper. The left row represents the PDB ids of the WT and mutants.

Information centrality indicates the role of residues in transmitting information across the network (29). Slight variations in the distributions of both 1e1w and 2k1d relative to 1qm2 suggest alterations in residual interactions.

### Identifying key residues through a combination of diverse centrality measures

We analyse the combined effect (30) of different node-based centrality measures, as each offers a unique insight into the network structure (see Methods). Beyond individual centrality assessments, we implemented an integrated network analysis approach utilising normalised combined centrality scores to identify structurally critical residues across all protein variants. The combined centrality score was computed through a two-step normalization process: first, individual centrality measures (degree, betweenness, closeness, eigenvector, and information centrality) were rescaled in the range [0, 1] using min-max normalization to ensure equal weighting across metrics and eliminate scale-dependent biases. Subsequently, the normalised values were summed to generate unified composite scores. Residues exhibiting combined centrality scores exceeding 1.25 standard deviations above the mean were classified as structurally critical nodes and were selected for further study to explore their structural and functional roles.

We find that in spite of variable point mutations in prion protein structure, five residues Met206, Val209, Val210, Met213, Cys214) are shared by all the protein structures with consistently high composite centrality value as well as individual measure values. Structurally, all the residues majorly reside in the C-terminal domain of the protein, which spans approximately 125-228 residues. C-terminal domain composed of three alpha helices [H1 (144-154), H2 (173-194), H3 (200-228)] and two stranded β-sheet [(β1 (128– 131), β2 (161–164)]. More specifically, Tyr150 lies within Helix 1, while Pro158 occurs just after H1 in the loop preceding β2. Val161, together with Tyr162 and Tyr163, belongs to the β2 strand. Phe175, Cys179, Val180, Thr183, Ile184, and His187 all fall within Helix 2, with His187 near its less well-ordered C-terminal end. Helix 3 encompasses Met206, Glu207, Val209, Val210, Glu211, Met213, Cys214 (linked via disulfide to Cys179), Ile215, Gln217, and Tyr218. These residues significantly affect overall network connectivity, according to network theory.

### Shortest Path Alteration Fraction

We analyze the values along with the integrated network centrality measures across the WT and mutants. Some residues show a distinct pattern of structural organization centered on specific critical residues. These residues define protein stability and communication pathways and reside in the C-terminal globular domain. The network analyses of prion protein variants using SPAF revealed variations in residue connectivity patterns compared to WT 1qm2.

In the case of WT, the *T_x_* is maximum for Val 209, came out to be a dominant hub and shows highly centralised nature in the network. A similar kind of network connectivity is also found in 1e1w, where Val209 is a part of 156 shortest paths. His187 exhibits varying *T_x_*, showed highest connectivity 1e1w (*T_x_* = 128), 1e1p (*T_x_* = 108), and 1qm2 (*T_x_* = 97) while comparatively reduced connectivity in 2k1d (*T_x_* = 63). Further, Tyr 218 showed the most striking variations from being a part of the highest number of shortest paths in 1e1g, where this residue is a part of almost 184 shortest paths and moderate variations in 1e1p *T_x_* = 53) and WT *T_x_* = 53). Another dominant hub is Phe175, in the case of 1e1p (*T_x_*= 101) and 1fkc (*T_x_* = 91), shares similarity with the WT 1qm2 (*T_x_* = 83), while showed moderate levels in the other variants. In the case of huPrP (Asp178Asn) mutant, the hyper centralised residue was Ile184 (*T_x_*= 117), while maintaining moderate levels in huPrP (Met166Val) (*T_x_*= 76), 1e1w (*T_x_* = 60), and 1qm2 (*T_x_* = 67). In 1e1p and 1fkc, 217Glu exhibited elevated *T_x_* levels of 111 and 84, respectively, as compared to the WT (*T_x_* = 54).

### Differential networks

The minor differences in network topologies observed in prion protein mutants reveal fundamental communication pathways that determine the molecular mechanisms/structural changes underlying pathogenicity. The dramatic shift in the *T_a_* value across the mutants reveals distinct hypercentralized residues and network architectures. The difference in closeness and information centrality distributions, along with other centrality measures, illustrate how mutations influence the network’s structural pattern. To analyze connectivity differences, we employ a DN approach, using node-based network centrality metrics, such as betweenness and closeness.

A substitution mutation at the 166^th^ position (Met166Val), has not been considered a lethal mutation. Although this residue sits in a very critical region of β2–α2 loop, where it links the β-sheet to the α2 helix. Using our DN analysis, we find that edges (Ile139, Met213), (Met205, Val210), (Tyr218, Tyr225), (Gln186, Thr191), (Met166, Tyr218), (Phe141, Tyr157), and (Tyr163, Cys214) show the largest changes in betweenness and closeness centrality upon removal. This change indicated that many shortest paths were rerouted in the mutant network.

Edge (Met166, Tyr218) links the β2–α2 loop to helix3 in the network, while in the case of WT huPrP, Tyr225 shares long-range interactions with Met166 and maintains the position of the β2–α2 loop (44). Aromatic and hydrophobic interactions among Met166, Phe175, Tyr218, and Tyr225 has also been affected by certain mutations such as., Pro212; these perturbations may increase the solvent exposure (45). Here, a large change in centralities after removing the edge (Met166, Tyr218) indicates that these residues could be a part of impactful communication pathways in the network.

Ser170Asn mutation in huPrP makes is more like Syrian hamster prion protein (shPrP). This mutation is present in a well-defined β2–α2 loop, although this loop is flexible in WT huPrP. NMR mutational studies reveal that a mutation at this site allows only restricted conformations (46). Here, we identified the following mutant-specific edges (Asp202, Glu207), (Met205, Val210), (Tyr163, Ile182), (His187, Lys194), (Ile182, His187), and (Arg156, Asn197). His187, Val180, and Tyr183 are the mutation sites strongly associated with the pathogenic conversion of cellular structure (47).

In the case of huPrp (Glu178Lys), from DN approach, identified edges are (His155, Glu196), (Ile138, Gln212), (Arg164, Asp167), (Tyr162, Arg164), (Tyr162, Thr190), (Asn174, Cys179), (Arg164, Cys179), (Met205, Val210), (Ile215, Ser222), (Val176, Glu221), (Arg156, Lys194), and (Ser143, Arg208). NMR structural data revealed that the overall fold of this mutant is similar to that of the WT, with altered electrostatic interactions around residue Lys200 (48). Stabilization of H2 and H3 depends on strong salt bridge interactions involving residues His155, Arg156, and Glu196. The substitution mutation at the 196th position with lysine breaks the interaction between Arg156 and Glu196 and destabilizes the hydrophobic interactions (49).

The Arg220Lys mutation shows how even conservative single amino acid substitutions in prion proteins can induce significant conformational changes. Despite both Arg and Lys being positively charged residues, the huPrP (Arg220Lys) variant exhibits profound structural deviations in helix 3 compared to huPrP (Met166Val), with high RMSD values across residues (Glu200, Tyr226). While helix-2 remains regular, helix 3 in this point mutant becomes notably irregular and displays a slight kink at residues (Tyr218, Glu220). (50). huPrP (R220K) mutant showed 58 newly formed edges after mutation, as compared to the WT huPrP (PDB: 1qm2). We analysed substantial changes in the global network topology after deleting edges one at a time in the parent network. The deletion of edges (Tyr162, His187), (His177, Glu207), (Met134, Lys220), and (Asn159, His187) significantly reduces overall betweenness centrality, thereby weakening long-range connections in the network. On the other hand, deleting (Asn159, His187), (Tyr162, His187), (His177, Glu207), and (Glu152, Arg156) significantly affected the network’s overall closeness centrality. (Met134, Lys220) edge established a connection between the N-terminal region near the β1-α1 loop and helix 3 of the C-terminal region. Similarly, the (Tyr162, His187) edge connects β2 and Helix 3.

After removing mutant-specific edges, we identified the node-wise changes in betweenness and closeness centrality by comparing them with the parent network. The residues most significantly affected are Tyr162, His187, Asn159, His177, and Glu207, indicating an important role in network communication when edges are disrupted. Disruption of (His177, Glu207) affects the inter-helical connection between helix 2 and helix 3, resulting in significant changes in the centrality measure. Furthermore, the removal of the (Asn159, His187) edge significantly increases the change in closeness centrality, which may enhance flexibility in the β1–α1 loop region. Together, these results indicate that Arg220Lys specific edges establish robust long-range communication pathways that are disproportionately disrupted, affecting a small set of key residues, likely stabilizing misfolded intermediates and promoting pathogenic conversion in prion diseases (50).

Another lethal and disease-associated mutation is huPrP (Asp178Asn). This mutation is important to study because it results in different phenotypes depending on the polymorphism at position 129. If the huPrP (Asp178Asn) mutation is present with Met129, it is linked to fatal familial insomnia, whereas the presence of Val at the 129^th^ position is associated with CJD (51). The most intriguing impact of this mutation is the disruption of the salt bridge between two highly conserved residues (Arg164, Asp178) (52). This alters the long-range interactions with Val129. Due to this point mutation, the removal of edges (Ile215, Ser222), (Val176, Glu221), (Arg156, Lys194), (Tyr162, Thr190), and (Ser143, Arg208) resulted in the maximum alteration in both overall betweenness (*B*) and closeness (*C*). An MD simulation study of the Arg208His mutation showed an association between this residue and conformational stability. It is clear that a mutation at Arg208 disrupts the salt bridge interaction within the network; for example, the Arg208Ala mutation eliminates the interaction between Asp144 and Arg208 (52). Further, Thr183Ala mutation removes two H-bonds between helix2 and beta sheet. Thr183 forms one H bond with Cys179, and another with Tyr162 (53). Altered interactions involving residues like Ile215, Ser222, Glu221, and Tyr225, which reside in helix 3, near the C-terminal region, become key determinants of conversion-prone forms (2).

### Secondary structure and thermal unfolding monitored by far-UV CD

Out of the five predicted residues, Met206, Val209, Val210, Met213, Cys214, we mutated Val210, Met213, and Cys214 with alanine by SDM. Two more residues, Gly126 and His140, were mutated in parallel to serve as negative controls. Far-UV CD spectra of WT prion protein and the four variants (Gly126Ala, His140Ala, Val210Ala, Met213Ala) were recorded between 200 and 250 nm over a temperature range of 30–90 °C. At 30 °C, all five proteins displayed the double minima near 208 and 222 nm characteristic of a predominantly α-helical fold, confirming that each variant refolded into a native-like conformation and that the substitutions. An increase in temperature induces a loss of helical signal at both minima and a shift of the spectral crossover, consistent with a transition from the folded α-helical state to a largely disordered ensemble.

Thermal transitions, plotted as the normalised change in ellipticity Δ[θ]/[θ], yielded melting temperatures of 70.8 °C for the WT, 70.1 °C for Gly126Ala, 68.7 °C for His140Ala, 64.2 °C for Val210Ala and 59.4 °C for Met213Ala. The variants therefore fall into two groups: Gly126Ala and His140Ala are essentially isostable with the WT (ΔT_m_ of −0.7 °C and −2.1 °C respectively), whereas Val210Ala and Met213Ala are substantially destabilised, with ΔT_m_ values of −6.6 °C and −11.4 °C. All proteins converged to a comparable residual ellipticity above 80 °C, indicating a similar unfolded end state.

### Molecular dynamics simulations of WT and mutant prion protein

MD simulations were carried out on the variants showing destabilising ΔΔG among the five predicted residues (Met206, Val209, Val210, Met213, Cys214), together with the two control substitutions (Gly126Ala, His140Ala). Gly126Ala, His140Ala, Met206Ala/Lys, Val209Lys, Val210Ala, Val210Lys, Met213Ala and Cys214Ala, each simulated alongside WT PrP for 500 ns.

WT PrP equilibrated rapidly and remained stable throughout, with a mean Cα RMSD of 0.18 nm and narrow fluctuations, indicating that the native fold is well maintained under the simulation conditions. The negative control substitution mutations broadly behaved similar to the WT. Gly126Ala, having an average RMSD of 0.18 nm, was indistinguishable from WT, and His140Ala showed only a marginal increase (0.23 nm). Predicted substitutions produced comparatively larger and more heterogeneous deviations. Val210Ala showed the highest mean RMSD (0.41 nm), rising progressively from ∼0.25 nm to a plateau near 0.5 nm between 200 and 450 ns, consistent with a sustained conformational rearrangement rather than a local fluctuation. Met213Ala (0.37 nm) and Val209Lys (0.36 nm) were comparably elevated, though with different trajectory shapes: Met213Ala sampled a high-RMSD state (∼0.4–0.6 nm) for the first 280 ns before dropping to a more stable ∼0.3 nm basin, whereas Val209Lys interconverted repeatedly between ∼0.2 nm and ∼0.45 nm states across the full trajectory, indicating persistent conformational exchange. Cys214Ala averaged 0.32 nm and was the only variant showing a clear upward drift in the final 50 ns, reaching ∼0.6 nm without re-equilibrating. Met206Ala (0.29 nm) and Val210Lys (0.25 nm) each showed a single high-RMSD episode — in the first ∼200 ns and around 250 ns, respectively, followed by relaxation to near-WT values for the remaining simulation.

RMSF profiles across residues 128–220 reproduced the expected pattern for the structured C-terminal domain in all systems, with minima over the helical segments and maxima at the loop regions and termini. Gly126Ala behaved almost exactly like WT, while the rest of the variants showed increased flexibility that was not confined to the mutation site but distributed across the domain, with the most pronounced increases in two regions: the 165–175 loop and the 190–200 segment preceding helix 3. Val210Ala showed the largest single peak (∼0.38 nm near residue 169), and Met213Ala showed broadly elevated fluctuation across residues 130–200 with a maximum near residue 196. His140Ala, Met206Lys and Val209Lys showed moderate, more localised increases, while Val210Lys remained close to WT over most of the sequence.

**Fig 2:**
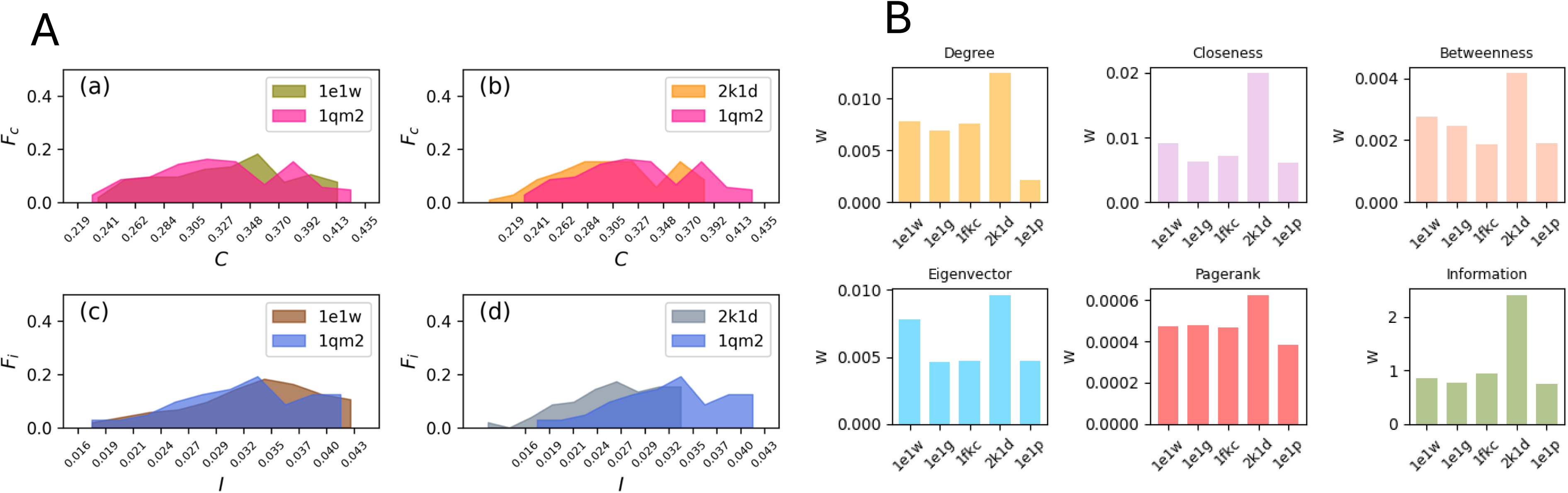
**[A]** Profile of how closeness centrality (a, b) and information centrality (c, d) is distributed for prion proteins 1e1w, 2k1d when compared with 1qm2. **[B]** Bar plots showing the Wasserstein distance (W) for six centrality measures (Degree, Closeness, Betweenness, Eigenvector, PageRank, and Information centrality) between the WT protein network (1qm2) and its mutants (1e1w, 1e1g, 1fkc, 2k1d, 1e1p). The Wasserstein distance quantifies how centrality distributions differ between the WT and each mutant, capturing topological shifts caused by point mutations. Among all measures, degree centrality shows the highest divergence, especially in the mutant 2k1d, thereby indicating significant local rewiring of network connectivity. Mutants 2k1d and 1e1w consistently display higher divergence across all measures, suggesting that even subtle point mutations can lead to substantial alterations in network architecture, impacting the connectivity and centrality of nodes.

**Fig 3:**
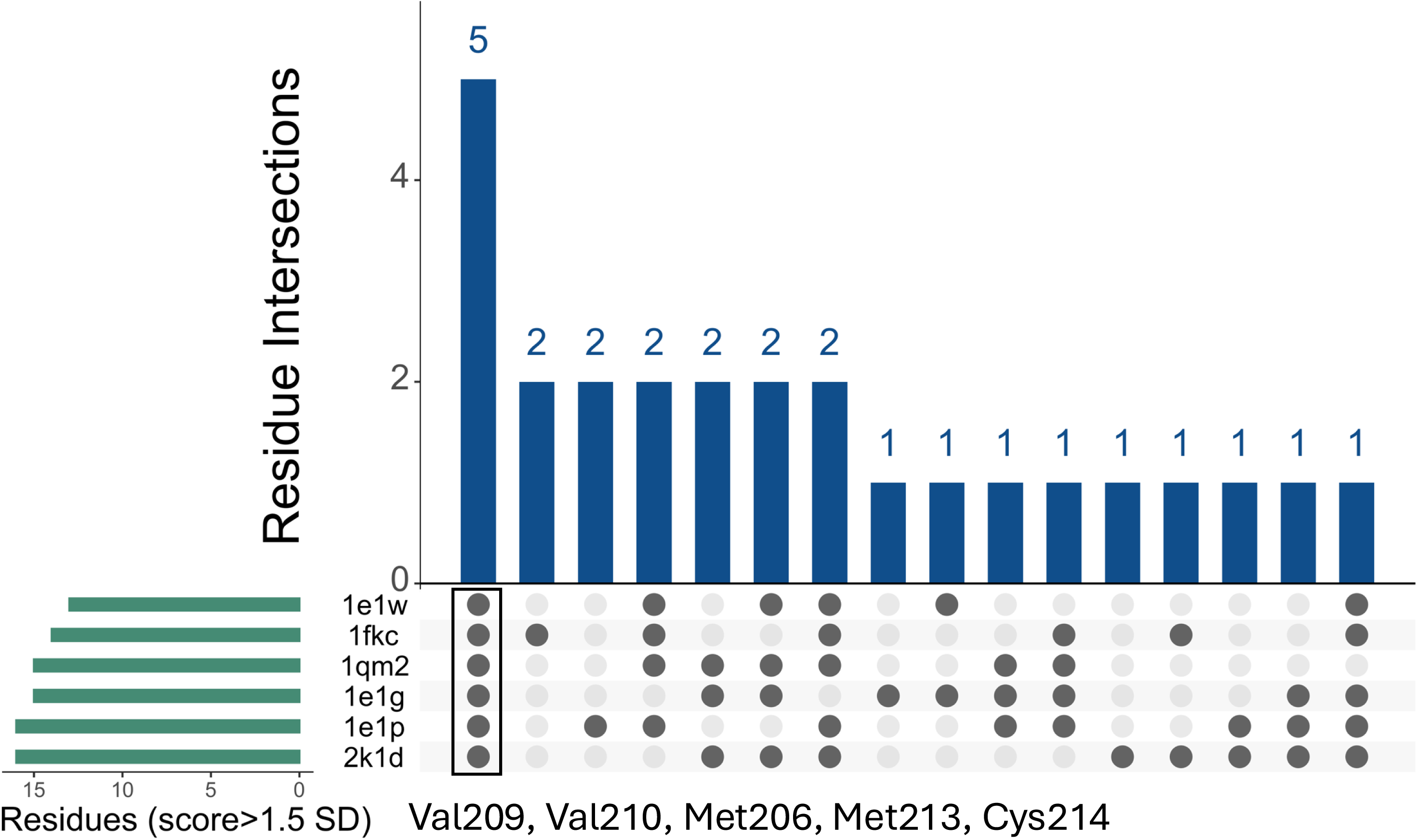
Analyses of protein residues bearing a score of network measure greater than 1.5 standard deviations across six different protein structures (PDB IDs: 1e1w, 1fkc, 1qm2, 1e1g, 1e1p, 2k1d). The blue bars indicate the number of common residues present in the protein structures. The highest intersection is observed for the combination Val209, Val210, Met206, Met213, and Cys214, which are shared by five structures. The matrix below the bar graph visually maps which residues are present in which structures, with filled circles indicating the presence of a residue in each protein. This visualisation highlights both conserved and unique residue patterns across the analysed proteins, suggesting potential hotspots of structural or functional significance.

**Fig 4:**
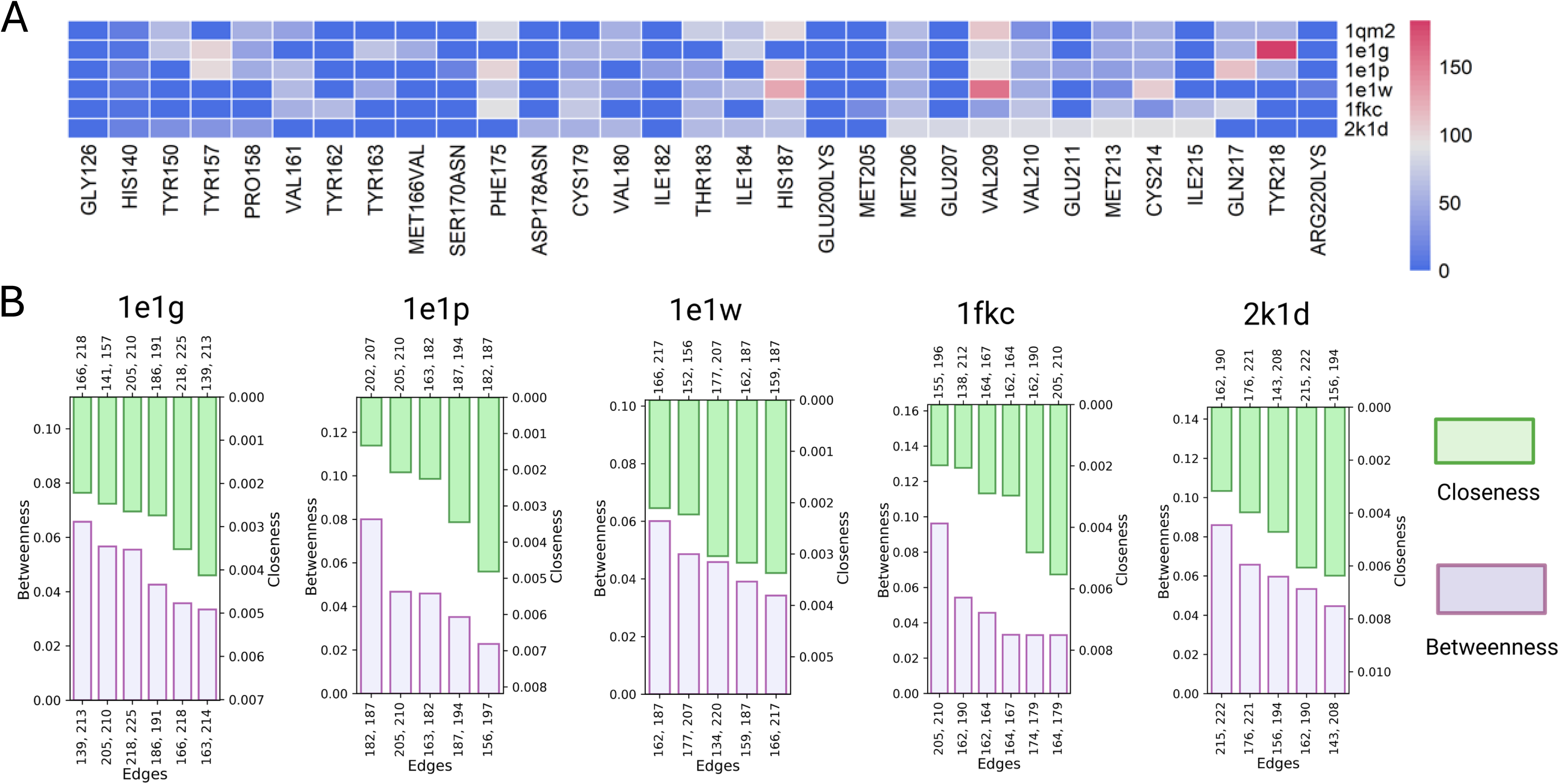
**[A]** Visualisation of SPAF analysis, where each cell displays the *T_x_* (SPAF=1), upon comparing the original/parent network and the new network, because of node removal. The x-axis denotes the specific residues targeted for removal. The y-axis contains PDB ids for the WT protein (1qm2) and the corresponding point mutant variant (2k1d, 1fkc, 1e1w, and 1e1p). The analyses compare the *T_x_* across different protein structures. Gly126 and His140 are the control residues, having a value below 1.25 σ threshold in the cumulative network centrality analysis for all the PDB IDs. **[B]** Key residue interactions found by differential network analyses. The effect of Betweenness centrality, (B), and Closeness centrality, (C), is significant on selected network nodes after removal of differential edges, which are unique in the mutants, when compared to the parent protein network (i.e., the network before edge deletion).

**Fig 5:**
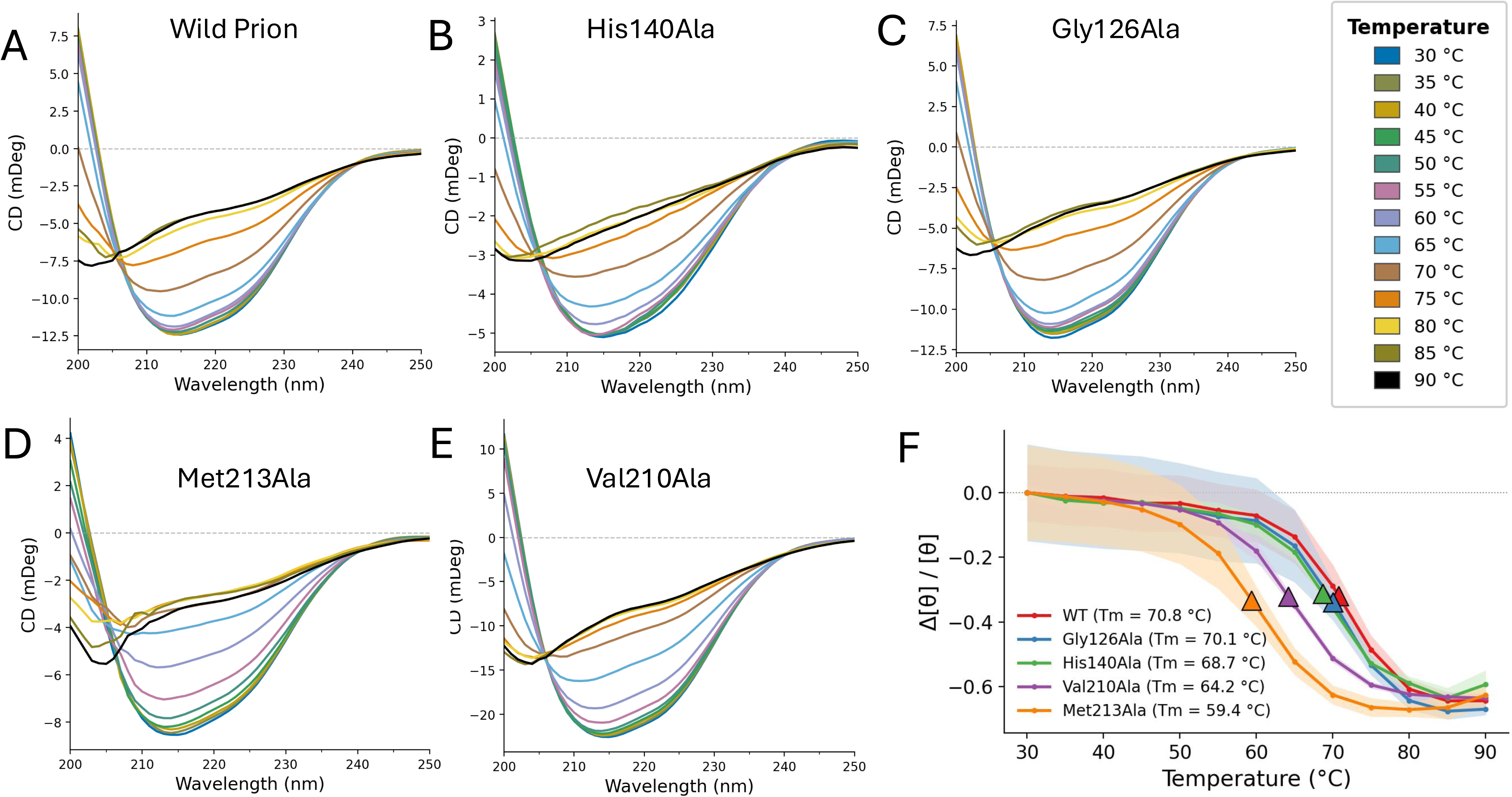
Far-UV CD spectra (190–260 nm) of WT and mutant proteins recorded at different temperatures. Thermal denaturation of WT (A) and mutants (B-E) monitored. Thermal unfolding was followed at 222 nm over the range [30–90] °C. Δθ/θ plotted against temperature for all variants, with the melting temperature T_m_ taken as the midpoint of the transition. The data points are the average of three experimental runs.

**Fig 6:**
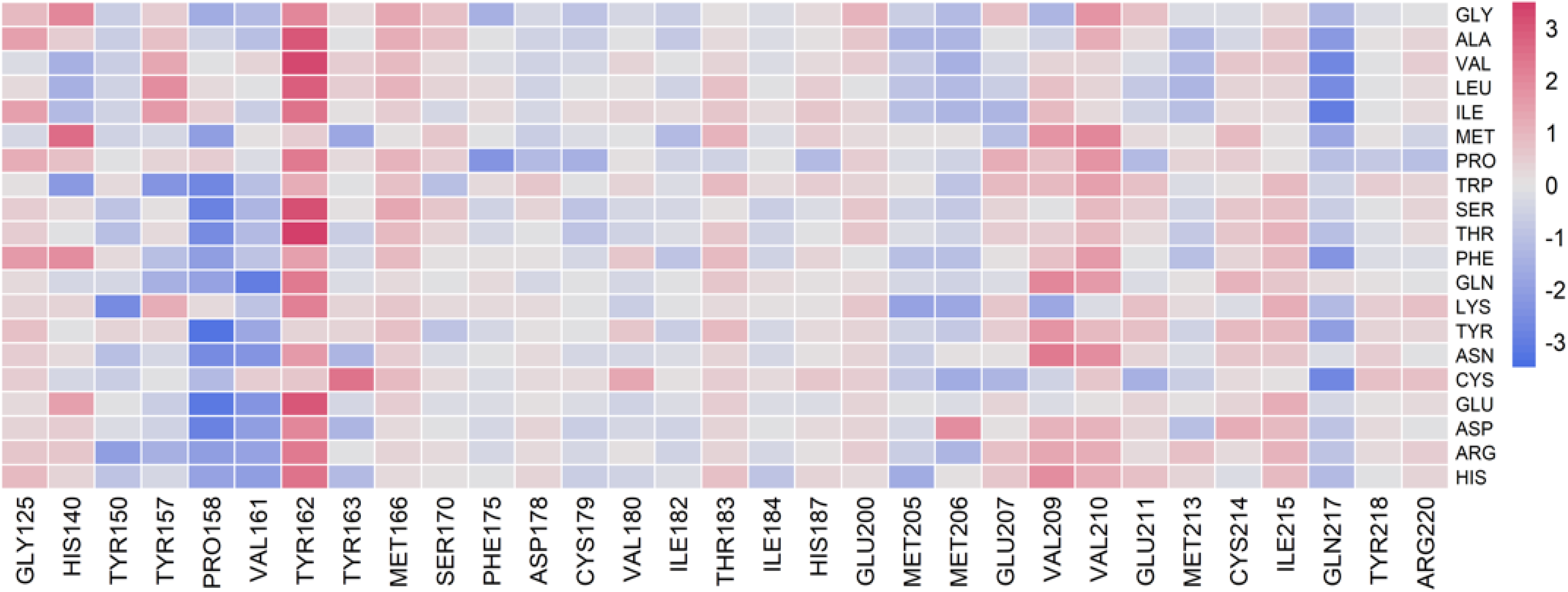
The protein mutant analysis heatmap, where the horizontal axis represents the residues having composite centrality scores exceeding 1.5 standard deviations above the mean, and the vertical axis represents all possible amino acid mutations at each selected residue position. Red indicates stabilising mutations that enhance WT protein stability (positive ΔΔG values), white represents neutral effects or no mutation (*ΔΔG* ≈ 0), and blue indicates destabilising mutations that reduce protein structural stability (negative ΔΔG values).

**Fig 7:**
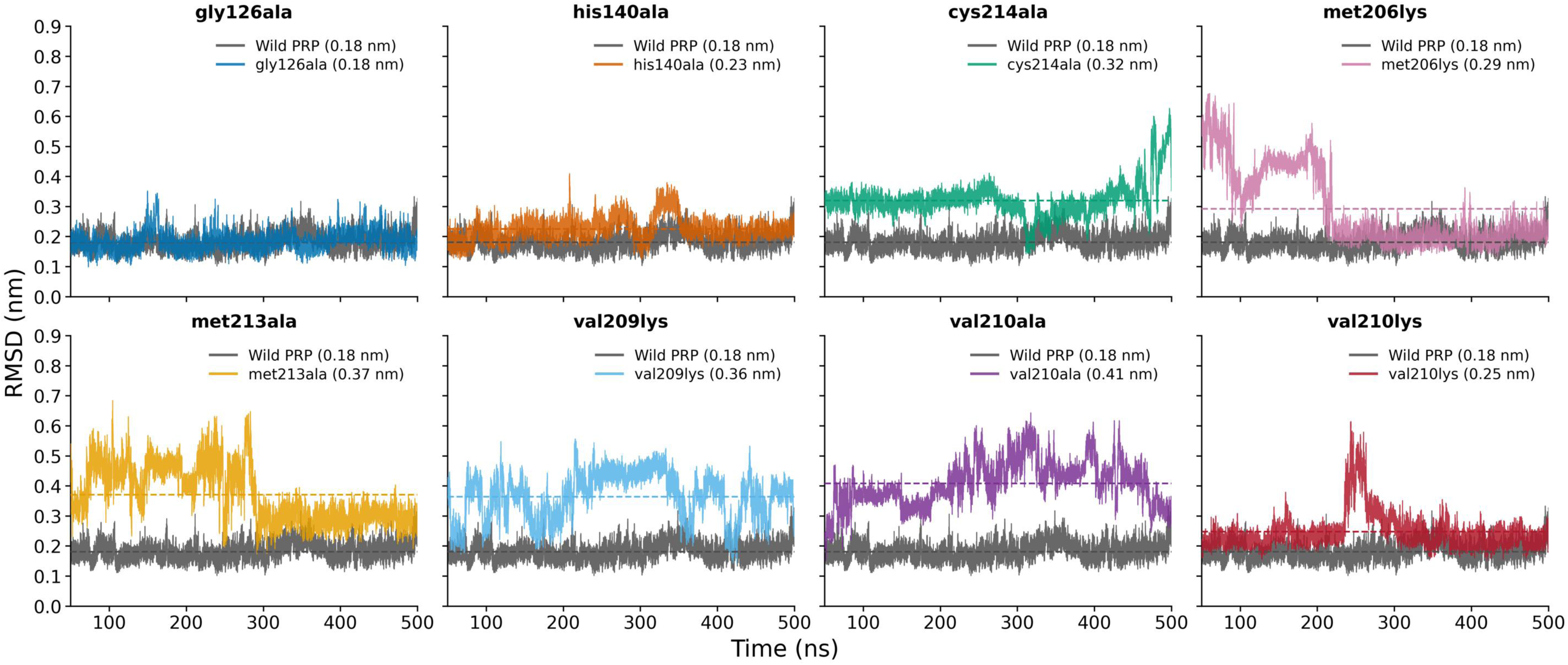
MD simulation of Backbone RMSD of WT and mutant huPrP(121-230) for 500 ns. Each panel demonstrates the WT overlaid with the mutant.

**Fig 8:**
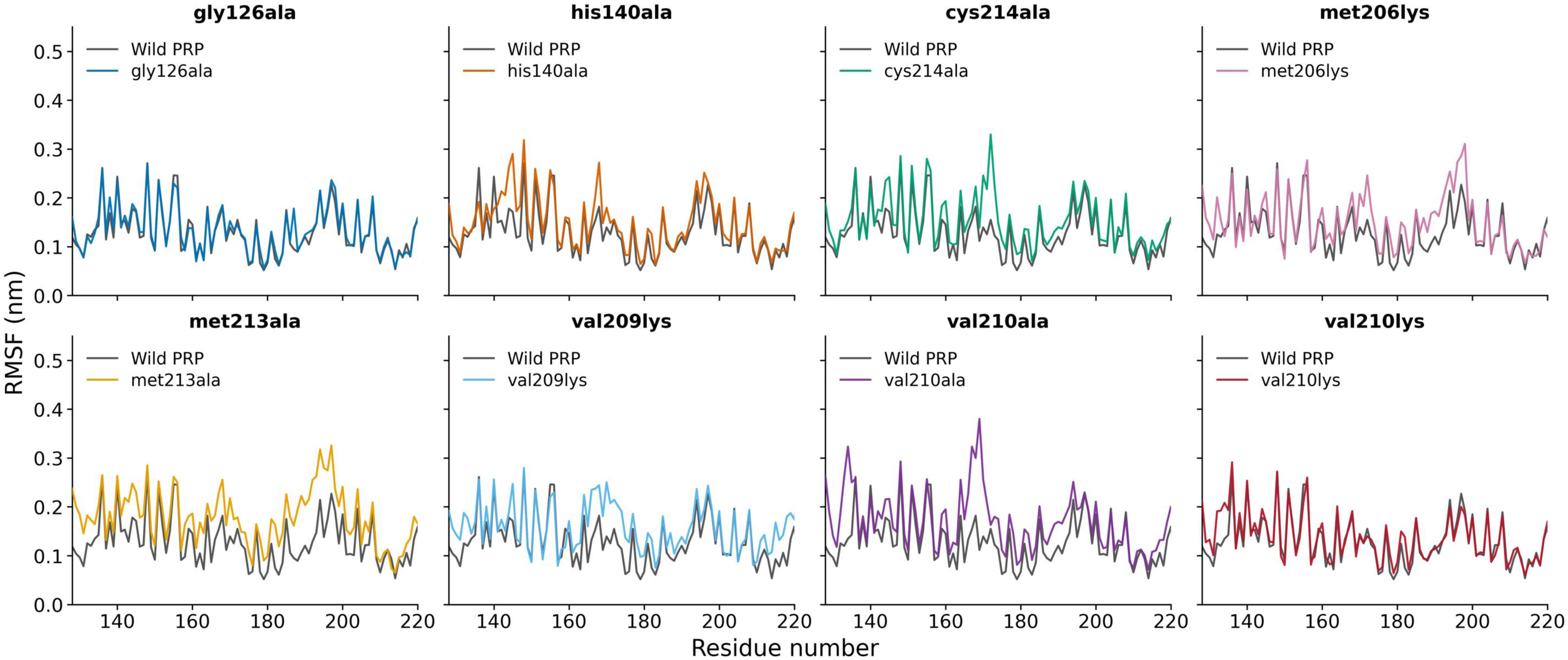
Per-residue C_α_ RMSF of WT and mutants huPrP(121-230). Each panel compares the WT with a mutant.

## Discussion and conclusion

Residue Interaction Networks provide a systems-level understanding of protein structures that is difficult to capture through sequence-or single-residue-based analyses (54, 55). In our study, we performed network analyses combined with a mutational analysis, providing integrated insight into protein stability. By employing a multiple centrality measures, coupled with SPAF analyses, and Differential Networks, we identified a small subset of residues that serve as critical hubs in both WT and mutant huPrP. This finding showed that the core packaging and long-range interactions of the network remained the same even after the point mutations. These residues are the central hubs of the network, primarily located in helices 2 and 3 (56), and serve as the initiation site for the conformational transition. The structural perturbations in these regions determine the propensity towards misfolding and aggregation of PrP^Sc^ (22, 56). Not only the common residues, but we also captured the shift in connectivity patterns in response to mutations through a DN approach. We employed the convergence of multiple independent computational metrics (57), which has the potential to provide robust validation of the biological significance of these identified hub residues. High SPAF values indicated that deleting these residues would disrupt and rewire the long-range communication paths throughout the network, thereby fundamentally altering inter-residue communication. Simultaneously, substitution mutations at these hub positions yielded large negative ΔΔG values, confirming substantial loss of protein stability (33).

Since network centrality and stability predictions based on ΔΔG are computational approaches, we next experimentally examine whether mutations at predicted residues affect structural stability.

### Structural and functional significance of the hub residues

Our findings of hub residues include Phe175 and Tyr169, which form a highly conserved aromatic-aromatic stacking interaction that maintains the orientation of the β2-α2 loop and overall protein stability (56). This region served as a critical conformational checkpoint because its conformation is determined by weak inter-residue interactions rather than a single dominant constraint. Mutations at Phe175 can produce extreme thermal destabilization. It has been observed that the Phe175Ala variant showed an 8°C decrease in unfolding temperature (58). This observation highlights how network-based approaches can correlate with experimentally observed “hot spots” where perturbations cause drastic effects on protein properties (59). Substitution mutation at Tyr169 not only affected the anchoring H-bond with Asp178 but also destabilized the aromatic interactions among Tyr169-Phe175-Tyr218. This destabilization enabled solvent-exposed rotamers of residue 169, thereby increasing the inter-helix distance and further destabilizing the α3 helix (60). Our simulations support this proposed mechanism directly: the variants with the largest loss of core packing showed their maximum backbone fluctuation at residue 169, indicating that the Tyr169 rotamer exposure inferred from structural arguments is reproduced as enhanced local mobility when the α2–α3 scaffold is weakened.

His187 is present at a crucial position, surrounded by charged residues Arg156, Glu196, and Asp202. This residue has a very low pKa value due to its buried microenvironment and is maintained within the α2 helix (61). His187Arg mutation is linked to familial CJD (62), introduced a permanent positive charge and disrupted the native charge balance within the helix. This charge replacement destabilized the electrostatic interactions between helix1 and helix2-3 and led to a beta sheet-rich pathogenic conformation. Hence, it acts as a pH-dependent switch that governs the structural stability of the entire helical bundle (63, 64).

Ile184, Met206, Val209/210, and Met213 stabilized inter-helical packing between α2 and α3 by participating in the formation of a hydrophobic scaffold (63). These highly connected residues stabilized both WT PrPᶜ and its mutants by providing multiple hydrophobic interactions, ensuring robustness even when individual interactions were disrupted. Mutations of the residues could catastrophically destabilize the inter-helical geometry and increase the propensity for the formation of the scrapie conformation (65). These hydrophobic core residues are highly conserved across mammalian species. This emphasized their structural relevance and suggested that the loss of these robust interactions would be evolutionarily disadvantageous. CD experiments on Met213Ala and Val210Ala showed a thermal destabilizing effect, with Val210Ala showing a ΔT_m_ of −6.6 °C and Met213Ala −11.4 °C. On the other hand, two negative control mutants remain within 2°C of WT. In MD simulations, Met206Lys and Val210Lys each sampled a destabilised state before returning to near WT RMSD values. In the case of Val210Ala and Met213Ala, packing defects were not recovered within 500ns.

CD can report the global stability of the folded protein but cannot specify the part of the protein, which is actually distorted. To check whether the network hubs behave as communication bottlenecks in a dynamic rather than a static structure, 500 ns simulations were performed on the variants that returned destabilising ΔΔG values among the predicted residues (Met206, Val209, Val210, Met213, Cys214), together with the two control substitutions for which CD data were obtained. WT PrP equilibrated rapidly at a mean Cα RMSD of 0.18 nm with narrow fluctuations, in agreement with the high structural stability of the folded domain reported in earlier simulations of human PrP (66, 67) and with its solution structures (9, 44).

Val210 lies within a hydrophobic core at the α2–α3 interhelical interface, forming direct hydrophobic contact with Val180. Substitution of Val210 with isoleucine is associated with CJD mutation. This reported mutation increases solvent exposure by reorienting several hydrophobic core residues and alters α2–α3 packing (42, 47). Met213 is partially buried, whereas neighbouring Met205 and Met206 are fully buried. Oxidation of the residue may disrupt hydrophobic interactions, thereby generating a molten globule with a high propensity for aggregation. This phenomenon is reported in the presence of another mutation, Glu200Lys, in which oxidation of Met213 is the initial step in the conformational change (68–70). A single native disulfide bond between Cys179 and Cys214 is the major determinant of tertiary structure. Absence or reduction of the bond may reduce equilibrium stability of both the native and aggregated states by approximately 3 kcal/mol (71–73).

### Connection of pathogenic mutations with hub residues

The destabilizing mechanism, involving pathogenic mutations Met166Val, Ser170Asn, Glu200Lys, Arg220Lys, and Asp178Asn, collectively converged to disrupt hydrophobic and electrostatic interactions with hub residues. M166V mutant is located at a flexible β2-α2 loop, showed a high cumulative centrality score as compared to other mutations. This mutation modified the hydrophobic environment and destabilized the stacking interaction between Phe175 and Tyr169 (59).

Mutations such as Asp178Asn and Glu200Lys altered the local charge density. Due to Asp178Asn (51) mutation, a negative charge is removed, which disrupts the essential interaction with somewhat exposed His177 within the α2 helix (62). Loss of a negative charge may destabilize the electrostatic interactions that anchor H2 and H3 in the structure. These local perturbations could increase the tendency of Prp^C^ to convert into a beta-sheet-rich scrapie form (PrP^Sc^). Another critical mutation that induced a positive charge in the α3 helix is Glu200Lys. The NMR study revealed that Glu200 is fully accessible to the solvent and is similar to the WT (1qm2). This substitution eliminated the salt-bridge interactions between Glu200 and Lys204, thereby altering the charge distribution on the protein surface (48).

Ser170Asn (33) and Asp178Asn are located at the junction of the α2-β2 loop. It has been observed that the mobility of the L1 loop (residues 166-175) and conformational dynamics are affected by the mutation of Ser170. Several hydrophobic interactions and H-bonds between the edges (Tyr128, Asp178) and (Tyr169, Asp178) are also affected by the mutation (52). Collectively, Ser170Asn and Asp178Asn mutations disturb the H-bonding with the residues that possess high centrality, such as Tyr169, Asp178, and Arg156. Furthermore, these interactions helped maintain the structure between the H2 and H3 helices by engaging aromatic amino acids Tyr150 and Tyr162. Met166Val (44) and Arg220Lys mutations affected H3 region. Specifically, Met166Val mutation affected the straightness of the entire helical axis.

In summary, our findings seem to indicate that the stability of the globular domain of huPrP is not dependent on an isolated residue or two, but rather on an interconnected network of residues. Regions involved in structural instability include (1) aromatic residues, (2) the electrostatic switch of His187, (3) perturbation in the α2–α3 hydrophobic scaffold, and (4) exposed hydrophobic surfaces. Mutating residues in these regions can alter the long-range interactions responsible for structural stability. Rigorous analyses by using three different techniques (centrality measures, shortest path alteration fraction, and differential networks) on Residue Interaction Networks, ΔΔG calculations, experimental studies through circular dichroism (CD) spectroscopy, and molecular dynamics simulations, independently highlight the effect of mutations. The strong agreement between these diverse methods augur well and helps in locating the destabilized regions that can initiate misfolding in related proteins. The broad framework demonstrated herein, presents wide applicability and should be of relevance in many other scenarios.

## References

1. Spagnolli G, Rigoli M, Orioli S, Sevillano AM, Faccioli P, Wille H, Biasini E, Requena JR. 2019. Full atomistic model of prion structure and conversion. PLOS Pathogens 15:e1007864.

2. Soto P, Gloeb GM, Tsuchida KA, Charles AA, Greenwood NM, Hendrickson H. 2023. Insight into the conserved structural dynamics of the C-terminus of mammal PrPC identifies structural core and possible structural role of pharmacological chaperones. Prion 17:55–66.

3. Van Der Kamp MW, Daggett V. 2009. The consequences of pathogenic mutations to the human prion protein. Protein Engineering, Design and Selection 22:461–468.

4. Zhou Z, Xiao G. 2013. Conformational conversion of prion protein in prion diseases. Acta Biochimica et Biophysica Sinica 45:465–476.

5. Deb A, Grewal RK, Roy S, Mitra D. 2020. Residue interaction dynamics in *Vaucheria* aureochrome1 light-oxygen-voltage: Bridging theory and experiments. Proteins 88:1660–1674.

6. Eghiaian F, Grosclaude J, Lesceu S, Debey P, Doublet B, Tréguer E, Rezaei H, Knossow M. 2004. Insight into the PrP^C^ → PrP^Sc^ conversion from the structures of antibody-bound ovine prion scrapie-susceptibility variants. Proceedings of the National Academy of Sciences 101:10254–10259.

7. Atkinson CJ, Zhang K, Munn AL, Wiegmans A, Wei MQ. 2016. Prion protein scrapie and the normal cellular prion protein. Prion 10:63–82.

8. Zahn R, Liu A, Lührs T, Riek R, Von Schroetter C, López García F, Billeter M, Calzolai L, Wider G, Wüthrich K. 2000. NMR solution structure of the human prion protein. Proceedings of the National Academy of Sciences 97:145–150.

9. Riek R, Hornemann S, Wider G, Billeter M, Glockshuber R, Wüthrich K. 1996. NMR structure of the mouse prion protein domain PrP(121–231). Nature 382:180–182.

10. James TL, Liu H, Ulyanov NB, Farr-Jones S, Zhang H, Donne DG, Kaneko K, Groth D, Mehlhorn I, Prusiner SB, Cohen FE. 1997. Solution structure of a 142-residue recombinant prion protein corresponding to the infectious fragment of the scrapie isoform. Proceedings of the National Academy of Sciences 94:10086–10091.

11. Knowles TPJ, Zahn R. 2006. Enhanced Stability of Human Prion Proteins with Two Disulfide Bridges. Biophysical Journal 91:1494–1500.

12. Knaus KJ, Morillas M, Swietnicki W, Malone M, Surewicz WK, Yee VC. 2001. Crystal structure of the human prion protein reveals a mechanism for oligomerization. Nature Structural Biology 8:770–774.

13. Kaur Grewal R, Mitra D, Roy S. 2015. Mapping networks of light–dark transition in LOV photoreceptors. Bioinformatics 31:3608–3616.

14. Sinha S, Samaddar S, Das Gupta SK, Roy S. 2021. Network approach to mutagenesis sheds insight on phage resistance in mycobacteria. Bioinformatics 37:213–220.

15. Berman HM. 2000. The Protein Data Bank. Nucleic Acids Research 28:235–242.

16. Berman HM, Battistuz T, Bhat TN, Bluhm WF, Bourne PE, Burkhardt K, Feng Z, Gilliland GL, Iype L, Jain S, Fagan P, Marvin J, Padilla D, Ravichandran V, Schneider B, Thanki N, Weissig H, Westbrook JD, Zardecki C. 2002. The Protein Data Bank. Acta Crystallographica Section D Biological Crystallography 58:899–907.

17. Grewal R, Roy S. 2015. Modeling proteins as residue interaction networks. Protein & Peptide Letters 22:923–933.

18. Khamaru M, Nath D, Mitra D, Roy S. 2024. Assessing Combinatorial Diversity of Aureochrome Basic Leucine Zippers through Genome-Wide Screening. Cells Tissues Organs 213:133–146.

19. Grant CD. 2023. Advances in Biology. Volume 31st ed. Nova Science Publishers, Incorporated, New York.

20. Newman MEJ. 2001. Scientific collaboration networks. II. Shortest paths, weighted networks, and centrality. Physical Review E 64:016132.

21. Hu G, Yan W, Zhou J, Shen B. 2014. Residue interaction network analysis of Dronpa and a DNA clamp. Journal of Theoretical Biology 348:55–64.

22. Amitai G, Shemesh A, Sitbon E, Shklar M, Netanely D, Venger I, Pietrokovski S. 2004. Network Analysis of Protein Structures Identifies Functional Residues. Journal of Molecular Biology 344:1135–1146.

23. Bonacich P. 1972. Factoring and weighting approaches to status scores and clique identification. The Journal of Mathematical Sociology 2:113–120.

24. Jeong H, Mason SP, Barabási A-L, Oltvai ZN. 2001. Lethality and centrality in protein networks. Nature 411:41–42.

25. Maslov S, Sneppen K. 2002. Specificity and Stability in Topology of Protein Networks. Science 296:910–913.

26. Negre CFA, Morzan UN, Hendrickson HP, Pal R, Lisi GP, Loria JP, Rivalta I, Ho J, Batista VS. 2018. Eigenvector centrality for characterization of protein allosteric pathways. Proceedings of the National Academy of Sciences 115.

27. Foutch D, Pham B, Shen T. 2021. Protein conformational switch discerned via network centrality properties. Computational and Structural Biotechnology Journal 19:3599–3608.

28. Stephenson K, Zelen M. 1989. Rethinking centrality: Methods and examples. Social Networks 11:1–37.

29. Brandes U, Fleischer D. 2005. Centrality Measures Based on Current Flow, p. 533–544. In Diekert, V, Durand, B (eds.), STACS 2005. Springer Berlin Heidelberg, Berlin, Heidelberg.

30. Banerjee SJ, Sinha S, Roy S. 2015. Slow poisoning and destruction of networks: Edge proximity and its implications for biological and infrastructure networks. Physical Review E 91:022807.

31. Roy S, Filkov V. 2009. Strong associations between microbe phenotypes and their network architecture. Physical Review E 80:040902.

32. Filkov V, Saul ZM, Roy S, D’Souza RM, Devanbu PT. 2009. Modeling and verifying a broad array of network properties. Europhysics Letters 86:28003.

33. Parthiban V, Gromiha MM, Schomburg D. 2006. CUPSAT: prediction of protein stability upon point mutations. Nucleic Acids Research 34:W239–W242.

34. Zahn R, Von Schroetter C, Wüthrich K. 1997. Human prion proteins expressed in *Escherichia coli* and purified by high-affinity column refolding. FEBS Letters 417:400–404.

35. Swietnicki W, Petersen RB, Gambetti P, Surewicz WK. 1998. Familial Mutations and the Thermodynamic Stability of the Recombinant Human Prion Protein. Journal of Biological Chemistry 273:31048–31052.

36. Liemann S, Glockshuber R. 1999. Influence of Amino Acid Substitutions Related to Inherited Human Prion Diseases on the Thermodynamic Stability of the Cellular Prion Protein. Biochemistry 38:3258–3267.

37. Corsaro A, Thellung S, Russo C, Villa V, Arena S, D’Adamo MC, Paludi D, Rossi Principe D, Damonte G, Benatti U, Aceto A, Tagliavini F, Schettini G, Florio T. 2002. Expression in E. coli and purification of recombinant fragments of wild type and mutant human prion protein. Neurochemistry International 41:55–63.

38. Abraham MJ, Murtola T, Schulz R, Páll S, Smith JC, Hess B, Lindahl E. 2015. GROMACS: High performance molecular simulations through multi-level parallelism from laptops to supercomputers. SoftwareX 1–2:19–25.

39. Huang J, Rauscher S, Nawrocki G, Ran T, Feig M, De Groot BL, Grubmüller H, MacKerell AD. 2017. CHARMM36m: an improved force field for folded and intrinsically disordered proteins. Nature Methods 14:71–73.

40. Jorgensen WL, Chandrasekhar J, Madura JD, Impey RW, Klein ML. 1983. Comparison of simple potential functions for simulating liquid water. The Journal of Chemical Physics 79:926–935.

41. Bussi G, Donadio D, Parrinello M. 2007. Canonical sampling through velocity rescaling. The Journal of Chemical Physics 126:014101.

42. Parrinello M, Rahman A. 1981. Polymorphic transitions in single crystals: A new molecular dynamics method. Journal of Applied Physics 52:7182–7190.

43. Michaud-Agrawal N, Denning EJ, Woolf TB, Beckstein O. 2011. MDAnalysis: A toolkit for the analysis of molecular dynamics simulations. Journal of Computational Chemistry 32:2319–2327.

44. Ilc G, Giachin G, Jaremko M, Jaremko Ł, Benetti F, Plavec J, Zhukov I, Legname G. 2010. NMR Structure of the Human Prion Protein with the Pathological Q212P Mutation Reveals Unique Structural Features. PLoS ONE 5:e11715.

45. Giachin G, Biljan I, Ilc G, Plavec J, Legname G. 2013. Probing Early Misfolding Events in Prion Protein Mutants by NMR Spectroscopy. Molecules 18:9451–9476.

46. Calzolai L, Lysek DA, Güntert P, Von Schroetter C, Riek R, Zahn R, Wüthrich K. 2000. NMR structures of three single-residue variants of the human prion protein. PNAS 97:8340–8345.

47. Biljan I, Ilc G, Giachin G, Raspadori A, Zhukov I, Plavec J, Legname G. 2011. Toward the Molecular Basis of Inherited Prion Diseases: NMR Structure of the Human Prion Protein with V210I Mutation. Journal of Molecular Biology 412:660–673.

48. Zhang Y, Swietnicki W, Zagorski MG, Surewicz WK, Sönnichsen FD. 2000. Solution Structure of the E200K Variant of Human Prion Protein. Journal of Biological Chemistry 275:33650–33654.

49. Cheng C, Daggett V. 2014. Molecular Dynamics Simulations Capture the Misfolding of the Bovine Prion Protein at Acidic pH. Biomolecules 4:181–201.

50. Calzolai L, Lysek DA, Güntert P, Von Schroetter C, Riek R, Zahn R, Wüthrich K. 2000. NMR structures of three single-residue variants of the human prion protein. 97:8340–8345.

51. Apetri AC, Vanik DL, Surewicz WK. 2005. Polymorphism at Residue 129 Modulates the Conformational Conversion of the D178N Variant of Human Prion Protein 90−231. Biochemistry 44:15880–15888.

52. Bamdad K, Naderi-manesh H, Baumgaertner A. 2013. Conformational instability of human prion protein upon residue modification: a molecular dynamics simulation study. EXCLI Journal 3:212–222.

53. Riek R, Wider G, Billeter M, Hornemann S, Glockshuber R, Wüthrich K. 1998. Prion protein NMR structure and familial human spongiform encephalopathies. Proceedings of the National Academy of Sciences 95:11667–11672.

54. Yehorova D, Di Geronimo B, Robinson M, Kasson PM, Kamerlin SCL. 2024. Using residue interaction networks to understand protein function and evolution and to engineer new proteins. Current Opinion in Structural Biology 89:102922.

55. Di Paola L, Giuliani A. 2015. Protein contact network topology: a natural language for allostery. Current Opinion in Structural Biology 31:43–48.

56. Chen J, Thirumalai D. 2013. Helices 2 and 3 Are the Initiation Sites in the PrP *^C^* → PrP *^SC^* Transition. Biochemistry 52:310–319.

57. Jalili M, Salehzadeh-Yazdi A, Gupta S, Wolkenhauer O, Yaghmaie M, Resendis-Antonio O, Alimoghaddam K. 2016. Evolution of Centrality Measurements for the Detection of Essential Proteins in Biological Networks. Frontiers in Physiology 7:375.

58. Christen B, Hornemann S, Damberger FF, Wüthrich K. 2012. Prion Protein mPrP[F175A](121– 231): Structure and Stability in Solution. Journal of Molecular Biology 423:496–502.

59. Huang D, Caflisch A. 2015. Evolutionary Conserved Tyr169 Stabilizes the β2-α2 Loop of the Prion Protein. Journal of the American Chemical Society 137:2948–2957.

60. Leske H, Hornemann S, Herrmann US, Zhu C, Dametto P, Li B, Laferriere F, Polymenidou M, Pelczar P, Reimann RR, Schwarz P, Rushing EJ, Wüthrich K, Aguzzi A. 2017. Protease resistance of infectious prions is suppressed by removal of a single atom in the cellular prion protein. PLoS ONE 12:e0170503.

61. Langella E, Improta R, Crescenzi O, Barone V. 2006. Assessing the acid–base and conformational properties of histidine residues in human prion protein (125–228) by means of pK *_a_* calculations and molecular dynamics simulations. Proteins 64:167–177.

62. Van Der Kamp MW, Daggett V. 2010. Influence of pH on the Human Prion Protein: Insights into the Early Steps of Misfolding. Biophysical Journal 99:2289–2298.

63. Malevanets A, Chong PA, Hansen DF, Rizk P, Sun Y, Lin H, Muhandiram R, Chakrabartty A, Kay LE, Forman-Kay JD, Wodak SJ. 2017. Interplay of buried histidine protonation and protein stability in prion misfolding. Scientific Reports 7:882.

64. Hosszu LLP, Tattum MH, Jones S, Trevitt CR, Wells MA, Waltho JP, Collinge J, Jackson GS, Clarke AR. 2010. The H187R Mutation of the Human Prion Protein Induces Conversion of Recombinant Prion Protein to the PrP^Sc^-like Form. Biochemistry 49:8729–8738.

65. Van Der Kamp MW, Daggett V. 2010. Pathogenic Mutations in the Hydrophobic Core of the Human Prion Protein Can Promote Structural Instability and Misfolding. Journal of Molecular Biology 404:732–748.

66. Gsponer J, Ferrara P, Caflisch A. 2001. Flexibility of the murine prion protein and its Asp178Asn mutant investigated by molecular dynamics simulations. Journal of Molecular Graphics and Modelling 20:169–182.

67. Zuegg J, Gready JE. 1999. Molecular Dynamics Simulations of Human Prion Protein: Importance of Correct Treatment of Electrostatic Interactions. Biochemistry 38:13862–13876.

68. Wolschner C, Giese A, Kretzschmar HA, Huber R, Moroder L, Budisa N. 2009. Design of anti- and pro-aggregation variants to assess the effects of methionine oxidation in human prion protein. Proceedings of the National Academy of Sciences 106:7756–7761.

69. Colombo G, Meli M, Morra G, Gabizon R, Gasset M. 2009. Methionine Sulfoxides on Prion Protein Helix-3 Switch on the α-Fold Destabilization Required for Conversion. PLoS ONE 4:e4296.

70. Wang Z, Feng B, Xiao G, Zhou Z. 2016. Roles of methionine oxidation in E200K prion protein misfolding. Biochimica et Biophysica Acta (BBA) - Proteins and Proteomics 1864:346–358.

71. Pastore A, Zagari A. 2007. A Structural Overview of the Vertebrate Prion Proteins. Prion 1:185–197.

72. Maiti NR, Surewicz WK. 2001. The Role of Disulfide Bridge in the Folding and Stability of the Recombinant Human Prion Protein. Journal of Biological Chemistry 276:2427–2431.

73. Honda R. 2018. Role of the Disulfide Bond in Prion Protein Amyloid Formation: A Thermodynamic and Kinetic Analysis. Biophysical Journal 114:885–892.

74. Gorfe AA, Caflisch A. 2007. Ser170 controls the conformational multiplicity of the loop 166– 175 in prion proteins: implication for conversion and species barrier. The FASEB Journal 21:3279–3287.

